# AVDM: Angular Velocity Decoding Model Accounting for Visually Guided Flight Behaviours of the Bee

**DOI:** 10.1101/654335

**Authors:** Huatian Wang, Qinbing Fu, Hongxin Wang, Paul Baxter, Jigen Peng, Shigang Yue

## Abstract

We present a new angular velocity estimation model for explaining the honeybee’s flight behaviours of tunnel centring and terrain following, capable of reproducing observations of the large independence to the spatial frequency and contrast of the gratings in visually guide flights of honeybees. The model combines both temporal and texture information to decode the angular velocity well. The angular velocity estimation of the model is little affected by the spatial frequency and contrast in synthetic grating experiments. The model is also tested behaviourally in Unity with the tunnel centring and terrain following paradigms. Together with the proposed angular velocity based control algorithms, the virtual bee navigates well in a patterned tunnel and can keep a certain distance from undulating ground with gratings in a series of controlled trials. The results coincide with both neuron spike recordings and behavioural path recordings of honeybees, demonstrating that the model can explain how visual motion is detected in the bee brain.

**Author summary:** Both behavioural and electro-physiological experiments indicate that honeybees can estimate the angular velocity of image motion in their retinas to control their flights, while the neural mechanism behind has not been fully understood. In this paper, we present a new model based on previous experiments and models aiming to reproduce similar behaviours as real honeybees in tunnel centring and terrain following simulations. The model shows a large spatial frequency independence which outperforms the previous model, and our model generally reproduces the wanted behaviours in simulations.

## Introduction

Insects, like flies and honeybees, though with tiny brains can deal with very complex visual flight tasks. It has been researched for decades how they detect visual motion. However, the neural mechanisms behind for explaining varieties of behaviours including patterned tunnel centring [1, 2] and terrain following [3–5] are still not very clear. According to the honeybees’ behavioural experiments performed, the key to their excellent flight control ability is the angular velocity estimation and regulation [6,7]. For instance, honeybees fly along the central path of narrow patterned tunnel carrying gratings of different spatial frequencies on both walls. The flight trajectory shifts towards the wall if the wall is moving along the fight direction, whilst away from the wall if it is moving in opposite direction, which indicate that honeybees adjust their positions by balancing the angular velocities estimated on both eyes [1]. Further electro-physiological experiments have also revealed that the spike responses of some descending neurons in honeybee’s ventral nerve cord grow as the angular velocity of the stimulus grating movement increases [8, 9], and the responses are largely insensitive to spatial frequency. Both indicate that honeybees can estimate the angular velocity of grating movement independently of its contrast or spatial frequency.

The angular velocity here is defined by the angular displacement Δ*ϕ* of the image motion in a small time interval Δ*t*, that is 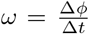. In tunnel centring and terrain following scenarios, denoting *v*_*x*_ as the forward flight speed and *d* as the distance to the surface, the angular velocity of image motion perceived by retina can also be expressed as 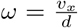. If the forward speed is maintained by a suitable constant forward thrust, then the distance to the surface will change automatically either by balancing the lateral angular velocities on both sides in tunnel centring, or by regulating the ventral angular velocity to a constant value in terrain following. Problems therefore arise as how honeybees estimate the angular velocity and further regulate it.

Due to the limited computation resources provided by the tiny brain, traditional computer vision methods, such as differential techniques, matching and feature-based approaches are restricted here [10]. Biological models usually correlate the light intensities of neighbouring photoreceptors with some temporal filters rather than calculate the spatial or temporal gradient of images [11]. Hassenstein and Reichardt [12] propose the first correlation motion detector which uses the temporal delay signal from left arm multiplies the non-delay signal from right to detect progressive motions (see Fig. 1(a)). A modified version, the HR-balanced detector (see Fig. 1(b)), consisting of two mirror-symmetrical subunits with a balance parameter is also proposed [13]. What’s more, the HR-detector based angular velocity sensor is designed [14] and has been applied to accomplish visual guided aircraft flight control tasks by Franceschini and Ruffier [15, 16]. However, both the HR model and the HR-balanced model are tuned for particular temporal frequency (number of gratings passed over the photoreceptor per second) rather than the angular velocity [13]. So the output of their sensor shows a large variance when tested by patterned ground in flight [14].

**Fig. 1.**
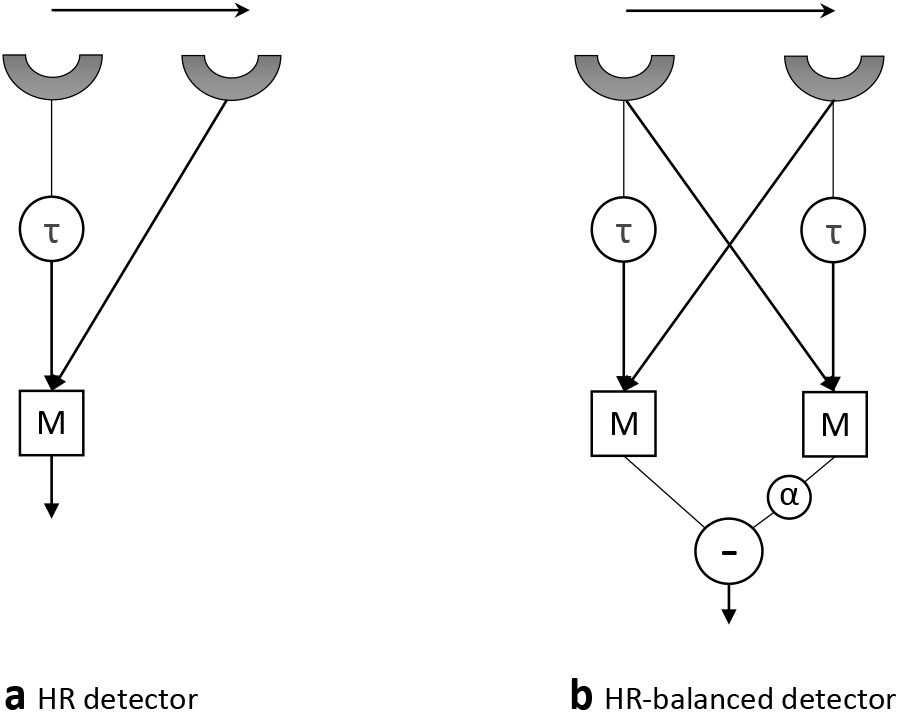
HR detector and HR-balanced detector. (a) In Hassenstein-Reichardt detector, a temporal delayed signal (*τ*) from left photoreceptor multiplies (M) the non-delay signal from right to give a preferred direction enhancement response [12]. (b) HR-balanced detector uses two subunits with a balance parameter (*α*) to tune the dependence on spatial frequency [13].

Based on their numerical analysis, Zanker et al. [13] suggest that the ratio of two HR-balanced detectors can produce a response tuned for angular velocity. Following this idea, Cope et al. [17] propose the C-HR model for estimating the angular velocity using the ratio of two HR-balanced detectors with different temporal delays. And the response is largely independent of the spatial frequency of the moving gratings especially when the angular velocity is around 100 °/*s*. However, this deviates from the biological experiments that honeybees usually fly at an angular velocity around 300 °/*s* [18]. Therefore the C-HR model can not fully match the aforementioned biological findings and explain the internal neural implementation of angular velocity estimation in the bee brain. Riabinina and Philippides [19] also build up a model, the R-HR model, using a channel fully dependent on temporal frequency as the denominator to get an angular velocity tuned response. The R-HR model performs well when the grating moving at a relatively low angular velocity. But the spatial independence gets weaker as the motion speed increases. Being inspired by the neural structure of Drosophila’s visual system, Wang et al. [20] propose a new motion detector with three inputs to produce a spatial independent response. Nevertheless, the model still does not show enough spatial independence for explaining the honeybee’s flight behaviours.

Moreover, previous mentioned three models [17,19,20] all use the ratio of two channels to tune a response for angular velocity, which may cause a problem of high output when the denominator is too small. It’s also one of the reasons why their models do not perform well when the angular velocity of the moving grating is low or high. This inspires us to build up a model without using the ratio of two channels, but to combine the spatial and temporal information from the moving gratings, based on the assumption that there are mechanisms combining both environmental texture and optic flow information in insect brains [21, 22].

Considering that insects’ compound eyes normally have thousands of ommatidia and a much higher temporal resolution than human, it’s possible to get the texture information from wide-field neurons and the temporal frequency information from spatially distributed HR-balanced detectors. Along with this idea, we propose an angular velocity decoding model (AVDM), and implement it into tunnel centring [23] and terrain following [24] simulations to reproduce similar behaviours of real bees. Under bio-plausible high sampling rate, we find the approximate square law relation between the model response and the temporal frequency when the temporal frequency is less than 50 Hz. By utilising this, the angular velocity decoding shows greater independence towards spatial frequency and contrast of the gratings than previous models. Series of controlled trails further verify the robustness of our model towards different conditions in visually guided flight simulations.

The rest of this paper is organised as follows. First the formulation of the model is presented and the control schemes for tunnel centring and terrain following are given. Then the results of synthetic grating experiments are exhibited to show the model’s independence to the spatial frequency and contrast of the grating. What’s more, the model is tested behaviourally by a virtual bee in tunnel centring and terrain following simulations. Finally we conclude and give some further discussions of this research.

## Materials and methods

### Input Signals Simulation

In order to explain the flight behaviours of honeybees well, the spatial and temporal resolutions of honeybees are investigated first to give a bio-plausible parameter setting. The ommatidia of bilateral compound eyes are arranged hexagonally separated by the interommatidial angle Δ*φ* (around 2°, varies in differnet regions [25]) and each corresponds to a visual column with an acceptance angle Δ*ρ* (about 2.5° [26]), as shown in Fig. 2. As for the temporal resolution, the critical fusion frequency (beyond which honeybees show no response to the flickering light source in electroretinogram test) is 165-300 Hz [27]. However, the behavioural experiments indicate honeybees can distinguish light fluctuation only when the stimuli is moving at temporal frequencies less than 200 Hz [28]. Therefore, we set the sampling rate as 200 Hz, in accordance with the honeybee’s ability of high temporal frequency image processing. Our proposed model is designed for dealing with this high sampling rate data and get a better performance in estimating angular velocity by using this high sampling rate data.

**Fig. 2.**
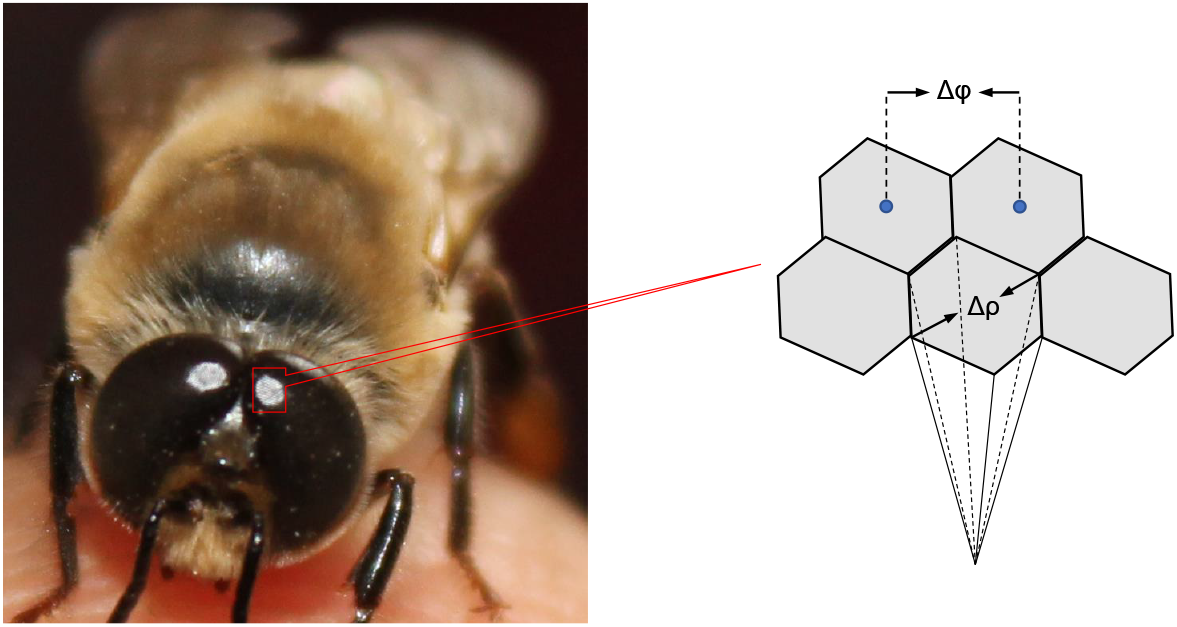
Illustration of the honeybee’s compound eye structures. The ommatidia are arranged hexagonally with an angular separation Δ*φ* (interommatidial angle) and each has its own small receptive field Δ*ρ* (acceptance angle).

The input signals are simulated as two dimensional image sequences of sinusoidal gratings moving across the field of view. Let λ and *ω* be the spatial period and the angular velocity of the grating movements, then the temporal frequency and angular frequency can be computed as *ω*/λ and 2*πω*/λ. Supposing the angular separation between pixels is *φ* (set to 2° in accordance with the honeybee’s spatial resolution), the input images can be expressed as following:

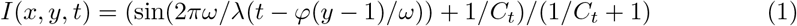

where (x,y) denotes the location of the ommatidium, t indicates the time and *C*_*t*_ ∈ (0, 1] is a parameter tuning the contrast of the gratings. Regarding our moving grating setting, *C*_*t*_ happens to be the image contrast at time t under Michelson contrast definition:

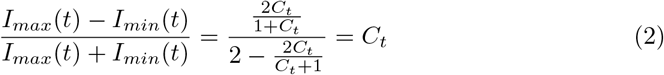

where the *I*_*max*_(*t*) and *I*_*min*_(*t*) (*I*_*max*_(*t*), *I*_*min*_(*t*) ≥ 0) indicate the highest and the lowest light intensities of the input signal at time t. For simplicity, normally we scale the pixel-wise intensity to be between 0 and 1 when the contrast is 1, except when considering the contrast invariance of the model.

### Angular Velocity Decoding Model

The model mainly contains three parts, the texture estimation pathway for spatial information extraction, the motion detection pathway for temporal information extraction and the decoding layer for angular velocity estimation. The structure of the proposed model is shown in Fig. 3. The global spatial frequency and image contrast information are estimated by the texture estimation part, and the motion information is processed by motion detectors. Then the angular velocity is decoded by combining both information.

**Fig 3.**
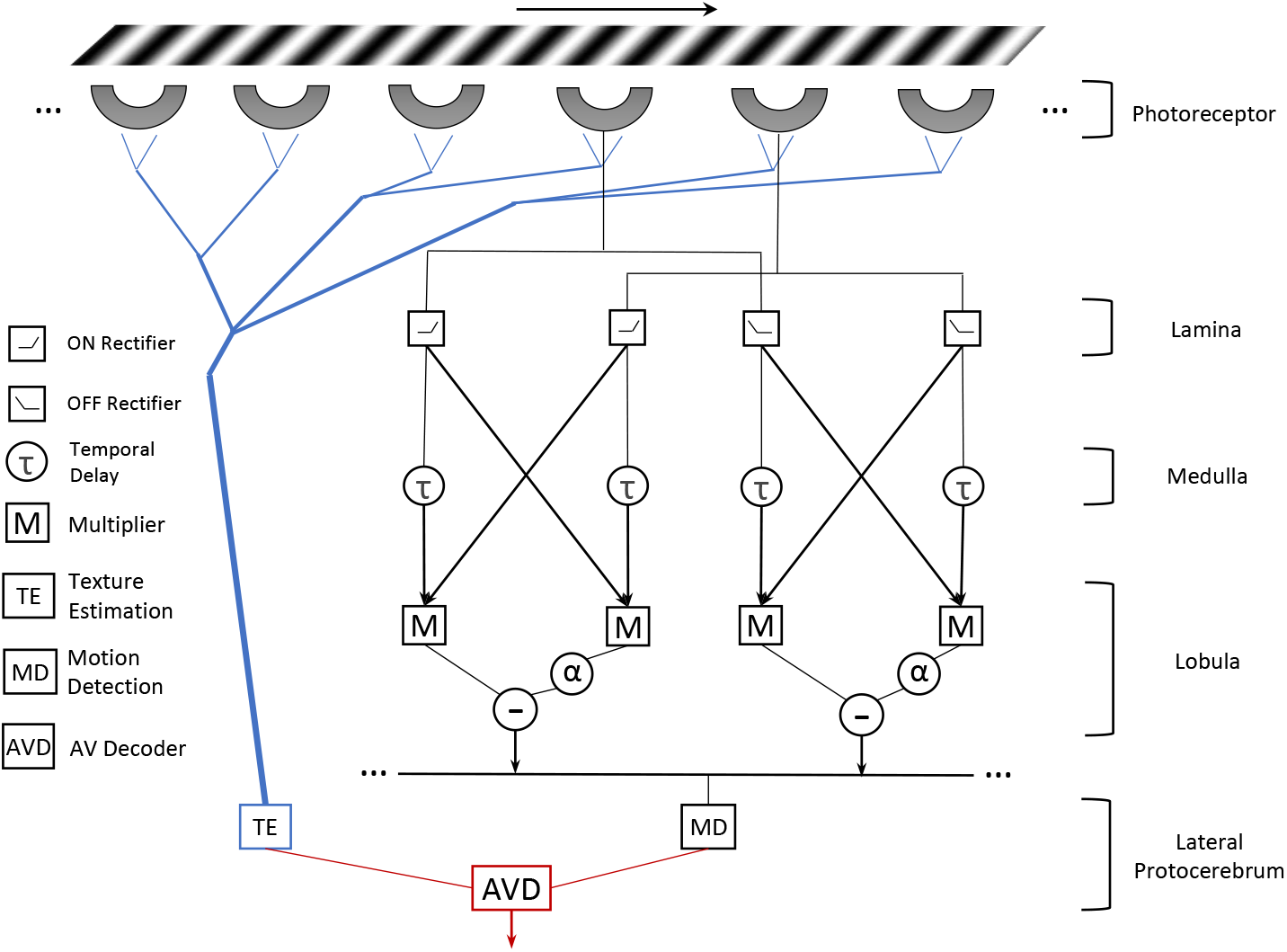
The structure of the proposed Angular Velocity Decoding Model. The visual information of grating’s moving is received by ommatidia. The texture information and the motion information across the whole vision field are combined in angular velocity decoding layer.

### Texture estimation pathway

The simulated input signals received by retina are first processed by the texture estimation pathway where the image contrast and the spatial frequency of the gratings are estimated by the light intensities of different locations. This is based on a hypothesis that insects can have a sense of the complexity of texture. Especially for the honeybees, they can discriminate patterns by visual cues including edge orientation, size and disruption [29], which indicates that the assumption is reasonable. Tunnel experiments also indicate that honeybees can distinguish the contrast on wall as low as 3%, and the flight speed in tunnel is little affect by the contrast provided that the contrast is larger than 3% [30]. One possible neural mechanism is to estimate the contrast and eliminate its affect in final response by the estimated value. Along this idea, the texture estimation pathway is proposed aiming to get the spatial frequency and image contrast with low computations.

Following the setting that every ommatidium covers 2° view [25], with 60 vertical(M) by 66 horizontal (N) receptors per eye covering the view of 120° by 132°, we can estimate the spatial period 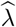 of the gratings according to the light intensities. First the input image is transferred into a binary image by the dynamic intensity threshold *I*_*thre*_ = (*I*_*max*_ − *I*_*min*_)/2. Then the spatial frequency is estimated by counting the number of the boundary lines of the binary image in the whole visual field. This simple method works well for sine-wave and square-wave gratings in our simulations. For more complex and detailed background, this method can also indicates the complexity of the textured background to some extent.

### Lamina layer

Insects interest the intensity change more than the intensity itself to detect motion. Therefore in our model, the input image frames are processed by the lamina layer where the light intensity change are computed to get the primary information of the visual motion [31]. Each photoreceptor computes the luminance change as the following:

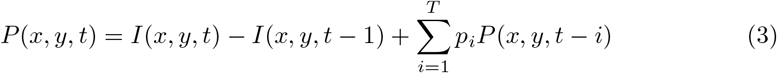

where *P* (*x, y, t*) corresponds to the luminance change of pixel (x,y) at time t; *T* denotes the maximum number of time steps the persistence of the luminance change can last and the persistence coefficients *p*_*i*_ ∈ (0, 1) is defined as

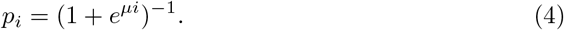

### ON and OFF layer

The luminance changes are separated into two pathways [32], the ON and the OFF pathways in this model. Specifically, the ON pathway deals with light intensity increments; whilst the OFF pathway processes brightness decrements. Denoting *f*^+^ = max(0, *f*) and *f*^−^ = min(0, *f*), then we can express the outputs of the cells in ON and OFF pathways as following:

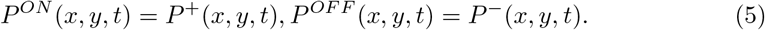

### Delay-and-correlation layer

Denoting *D*^*ON*^(*x, y, t*), *D*^*OFF*^(*x, y, t*) as the outputs of the ON and the OFF detectors and *τ* as the temporal delay in HR-balanced detectors, we have the following expression according to the structure of motion detectors in Fig. 3, where each pairwise neighbouring ON/OFF cells correlate with each other as:

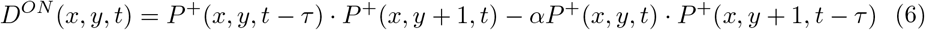

where *α* is chosen as 0.25 forming a partial balanced model [13]. And *D*^*OFF*^(*x, y, t*) can be expressed similarly.

### Angular velocity decoding layer

If the input signals are simulated using (1), we can get the outputs of the ON detectors by (6). The outputs of all ON and OFF detectors can be combined to get a response encoding the temporal frequency. In fact, the angular velocity of the background moving is caused by the insect’s ego-motion. Considering when a bee flying over a flat surface without changing its altitude, the consistency of the image motion speed helps us simplify the problem so that we can average the output signals from all ON or OFF detectors in visual field to get the final response *R*(*ω*, λ), which encodes the angular velocity:

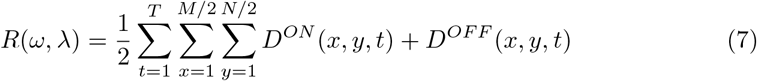

Here we take only one detector for example to analyse how the response is affected by the input signals. Let *S*_1_, *S*_2_ denote the luminance change of ommatidium A (left) and B (right), and 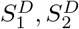 denote the temporal delayed luminance change of A and B, then according to the structure of HR-balanced detector (Fig. 1(b)), the response of the detector *R*_0_ can be expressed as 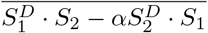, where the overline means the response is averaged over a time period to remove fluctuation caused by oscillatory input. For simplicity, we assume the image contrast is constant in this time period. Then the reponsonse *R*_0_ can be roughly expressed in theoretical terms [13] as the following equation:

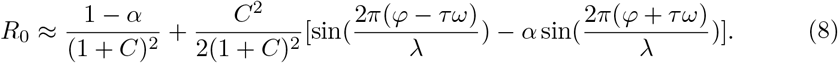

However, it is hard to derive angular velocity directly from (7) or (8). But by combining the texture information, we can decode the angular velocity information from the response *R*(*ω*, λ) using an approximation method. Though there is an inevitable fitting error, we can decrease it into an acceptable level if the fitting function is chosen well. The main idea is that using Taylor series omitting higher-degree Taylor polynomials to approximate (8), and using a power function of 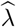 to tune the spatial independence. Along this idea, one decoding function can be chosen as following to approximate the actual angular velocity:

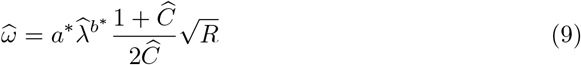

where 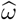 denotes the decoded angular velocity, 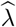 is the estimated spatial period and 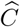 is the estimated contrast from texture estimation layer. Parameters *a*\* and *b*\* can be learned by minimizing the difference from the ground truth using alternate iteration method:

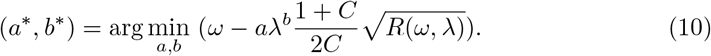

### Control Scheme for Tunnel Centring

By utilising the physics engine of Unity, the AVDM first has been embodied in a virtual bee to simulate the tunnel centring behaviour of the honeybee. Honeybee can centre itself in a narrow tunnel by balancing the angular velocities on both eyes [1]. Following this visual flight strategy, an AVDM-based control scheme is required to reproduce this behaviour. Then the performance of the model will be investigated by checking whether the virtual bee can centre itself in tunnel as real bees.

The control scheme for tunnel centring is described in Fig. 4. For simplicity, the forward flight speed is maintain the same, and we only focus on centring by the horizontal position controller using the difference between the angular velocities estimated by the AVDM on both eyes. Only lateral other than frontal vision field is considered in our simulation. We also assume that the orientation of head is roughly parallel to the central path of the tunnel and is seldom affected by the body movement. In fact, honeybees do perform a gaze stabilization in flight which use the head yaw turn ahead of the body yaw turn to against the orientation rotation [33].

**Fig 4.**
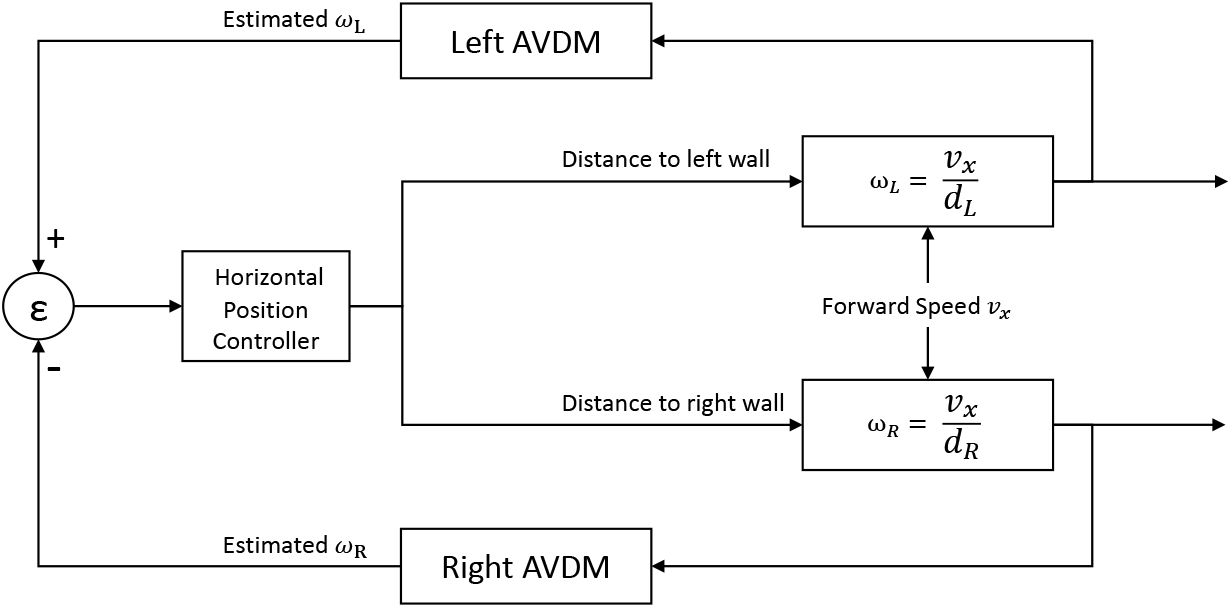
The AVDM-based closed loop control scheme for tunnel centring. The horizontal position controller is triggered by the difference *ɛ* between angular velocities estimated by left and right eyes.

Following the scheme, the virtual bee can adjust its position in tunnel automatically using only visual information in real time. The distance to left wall will increase if the difference *ɛ* is positive and vice versa. With little modification, the scheme can further be used to simulate the situations when one of the walls is moving along or against the flight direction.

### Control Scheme for Automatic Terrain Following

Honeybees will adjust their flight height to restore a preferred ventral angular velocity if the grating moves along the flight direction. This angular velocity regulating strategy helps honeybees navigate safely through tunnels [4]. In addition, the visual strategy is also used in aircraft automatic terrain following experiments [15]. The proposed model can be inspected in this flight task to see if it can improve the accuracy of the angular velocity regulation. Using the AVDM, we can estimate the angular velocity in flight. By regulating it accordingly to a constant value, the altitude will change automatically regardless of the prior knowledge to the exact altitude and forward flight speed.

The closed loop control scheme for terrain following referenced Franceschini’s work [15] is given in Fig 5. For simplicity, we assume the forward flight speed is maintained the same by a proper constant forward thrust. In fact, this assumption is reasonable since honeybees tend to adjust its flight height rather than speed to regulate the ventral angular velocity [4]. Thereby, the proposed AVDM can adjust the vertical lift according to the difference between the preset angular velocity and the consecutive estimated values. Here, the preset angular velocity is also estimated by the AVDM in the beginning phase when the vertical lift is set to the same value of gravity and where the ground is flat. After that, when the ventral angular velocity varies as a result of terrain undulating, the vertical lift controller will change the lift according to the difference *ɛ* between ventral angular velocity estimated and the preset value. If the difference *ɛ* is positive, the lift will increase and vice versa.

**Fig 5.**
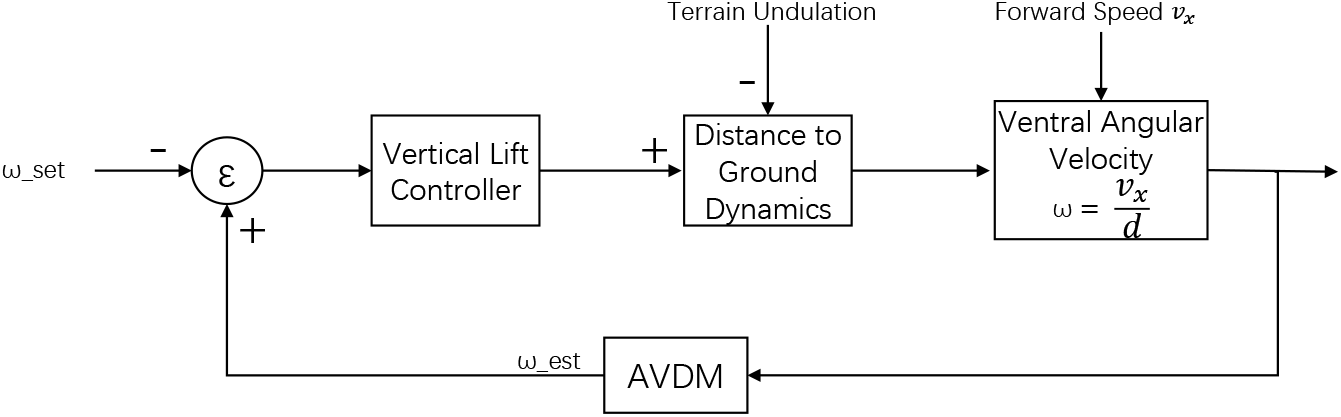
The AVDM-based closed loop control scheme for terrain following. The vertical lift controller is triggered by the difference *ɛ* between preset angular velocity *ω*_*set*_ and the estimated angular velocity *ω*_*est*_.

During the terrain following approach, the vertical speed *v*_*z*_ is relatively small, and the air resistance can be approximated as *f* = *kv*_*z*_. Then the vertical dynamics can be described using the following differential equations:

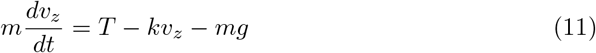

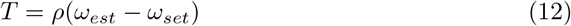

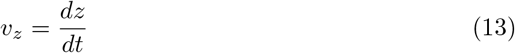

where m is the mass of the virtual bee, g is the gravity acceleration and T is the vertical lift. Given the initial conditions, then the flight trajectory can be computed step by step. In our simulation, this process can be achieved by the physics engine of Unity.

### Parameter Setting

Parameters of the proposed model and the control scheme are shown in **Table 1**. Parameter setting are mainly tuned manually based on empirical knowledge and stay the same in the following simulations unless particularly stated.

**Table 1.**
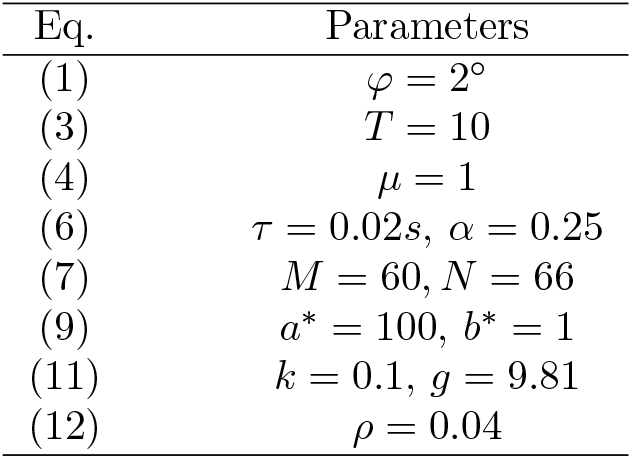
Parameters of the model and the control scheme

## Experiments and Results

Within this section, we present the experiments and results. The proposed model was first tested by synthetic grating stimuli to show its large spatial independence and robustness to contrast and noise in Matlab (© The MathWorks, Inc.). Then the model is implemented into a virtual bee by Unity (© Unity Technologies) to simulate the tunnel centring and terrain following behaviours of the honeybees.

### Angular velocity decoding results

To inspect the spatial frequency independence of the proposed model, sinusoidal gratings of a wide range of spatial periods (12° to 72°) are chosen as the visual inputs. The results of estimated angular velocities are shown in Fig. 6. The proposed model decodes the angular velocity well with little variance, except when narrow grating (12°) moving at a high angular velocity (larger than 700°/*s*). Honeybees tend to maintain a constant angular velocity of 300°/*s* [18], around which our proposed AVDM fits well and represents large spatial independence.

**Fig 6.**
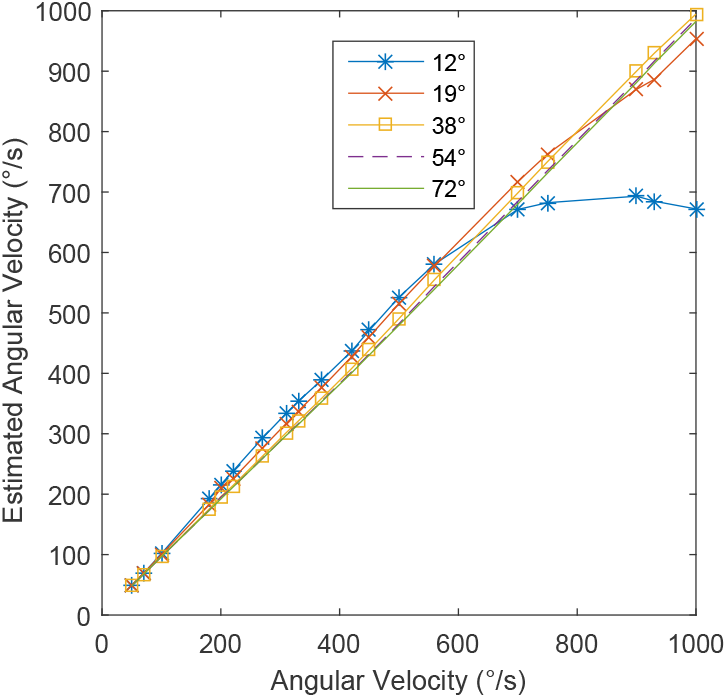
The angular velocity decoding results. The estimated angular velocity curves under different angular velocities when tested by moving gratings of different spatial periods (12°, 19°, 38°, 54° and 72°), demonstrating the spatial frequency independence of the model.

The adjusted R-squared values for different spatial periods are provide in **Table 2** to evaluate the biases of the decoded angular velocities from the ground truth. Most of the decoding curves estimate the angular velocity well since the adjusted R-squared values are close to 1. This means the AVDM performs stably to estimate the angular velocity against a wide range of spatial periods, explaining that honeybees navigate well in cluttered environment in an open flight.

**Table 2.** The adjusted R-squared values of angular velocity decoding curves of different spatial periods.

| Spatial Period | $12^\circ$ | $19^\circ$ | $38^\circ$ | $54^\circ$ | $72^\circ$ |
| --- | --- | --- | --- | --- | --- |
| Adjusted-$R^2$ | 0.8685 | 0.9962 | 0.9995 | 0.9981 | 0.9974 |

In order to demonstrate the improved spatial independence of the AVDM compared to the state-of-the-art angular velocity estimation models, we contrast it with two other aforementioned detecting models, the R-HR model [19] and the C-HR model [17]. The original results of their models are re-plotted in Fig. 7 under the same metric. In general, our model shows a much stronger spatial independence than the comparative models. The R-HR model shows a larger variance of responses when the angular velocity increases (see Fig. 7a). The C-HR model performs very well around 100°/*s*, but shows a larger variance at low (less than 60°/*s*) and high (faster than 300°/*s*) angular velocities. The normalized response lies in interval (0.8, 1) when the angular velocity is larger than 300°/*s*, which means it is hard for the C-HR model to discriminate approximate angular velocities in this region (see Fig. 7b). Compared to the comparative models, our proposed AVDM can better decode angular velocities with small variances.

**Fig. 7.**
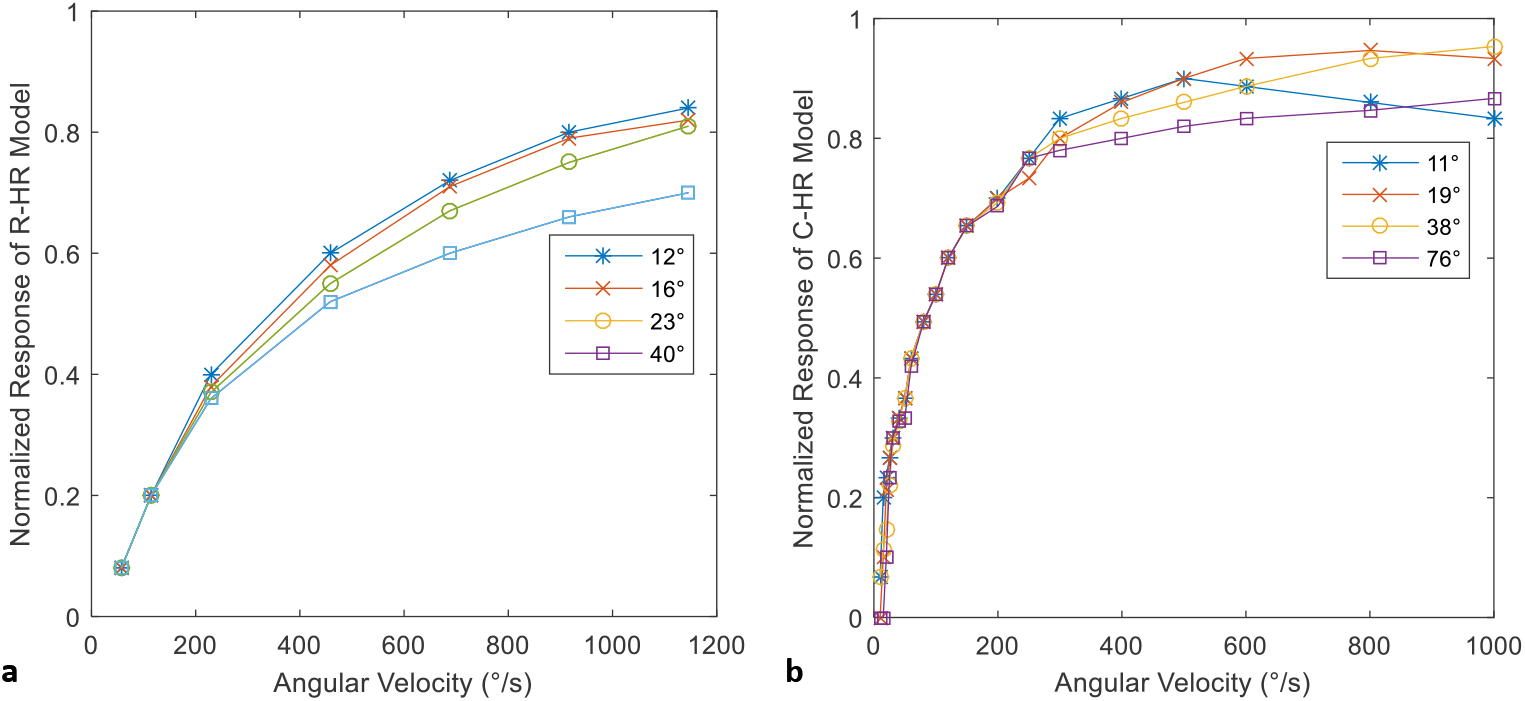
Contrast of responses of two other comparative models. (a) Response curves of the R-HR model for different spatial periods show roughly spatial independence [19]. (b) The C-HR model shows largely spatial independency around the velocity of 100 °/*s* [17].

### Image Contrast and Noise Invariance

Honeybees are able to navigate well in tunnel at varies of pattern contrasts on wall [30]. To evaluate the robustness of the model towards image contrast, we test the proposed model by the moving sinusoidal gratings of different contrasts. As can be seen from Fig. 8(a), the results show little variance when the image contrasts varies from 1/5 to 3/5. This outperforms previous model especially when the angular velocity is low and high [17]. The proposed model processes the texture estimation pathway where the contrast is first estimated and also the decoding layer where the estimated contrast is used to decode the angular velocity. The angular velocity is estimated accurately despite the contrast dynamics of input stimuli, reminiscent of honeybee’s flights through dynamic and cluttered environments.

**Fig. 8.**
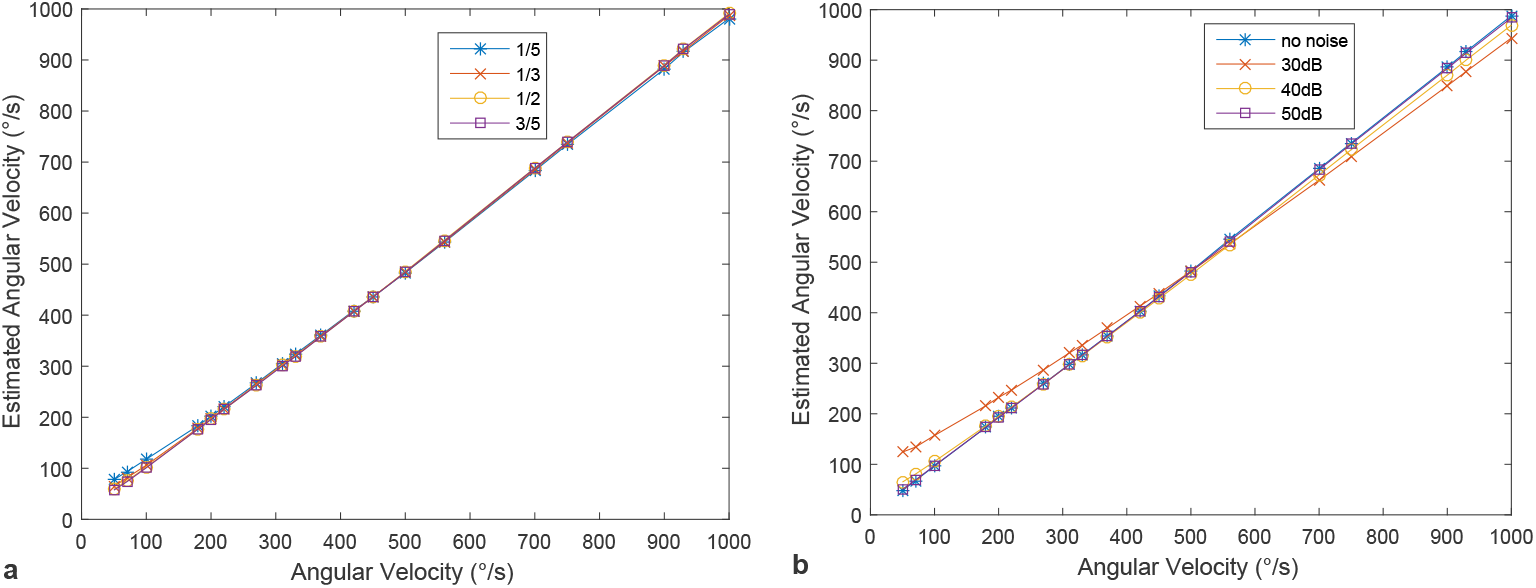
Robustness of the proposed AVDM against contrast and noise. (a) The proposed model is tested by the moving sinusoidal gratings of 54° spatial period and the image contrasts are set as 1/5, 1/3, 1/2 and 3/5. (b) The proposed model is tested by the moving sinusoidal gratings of 54° spatial period and the SNR of the input gratings are set as 30 dB, 40 dB and 50 dB. The result of the input without noise is also given as a reference.

To show the robustness against noise, we test the proposed model by adding different levels of Gaussian white noise into the input signals. The results can be seen from Fig. 8(b). The estimated image velocity curves show little variance when the signal-to-noise ratio (SNR) larger than 40 dB. The results demonstrate the reliability of the AVDM for estimating image velocity in an environment containing detailed background noise.

### Tunnel centring simulation results

To verify the effectiveness of the proposed AVDM in reproducing honeybees’ flight behaviours in narrow tunnel centring experiments, the virtual tunnel environment is set up with Unity (see Fig. 9), and a series of centring simulations have been performed. Using Unity engine, the images received by two eyes can be processed separately in real time to regulate the route of the flight.

**Fig. 9.**
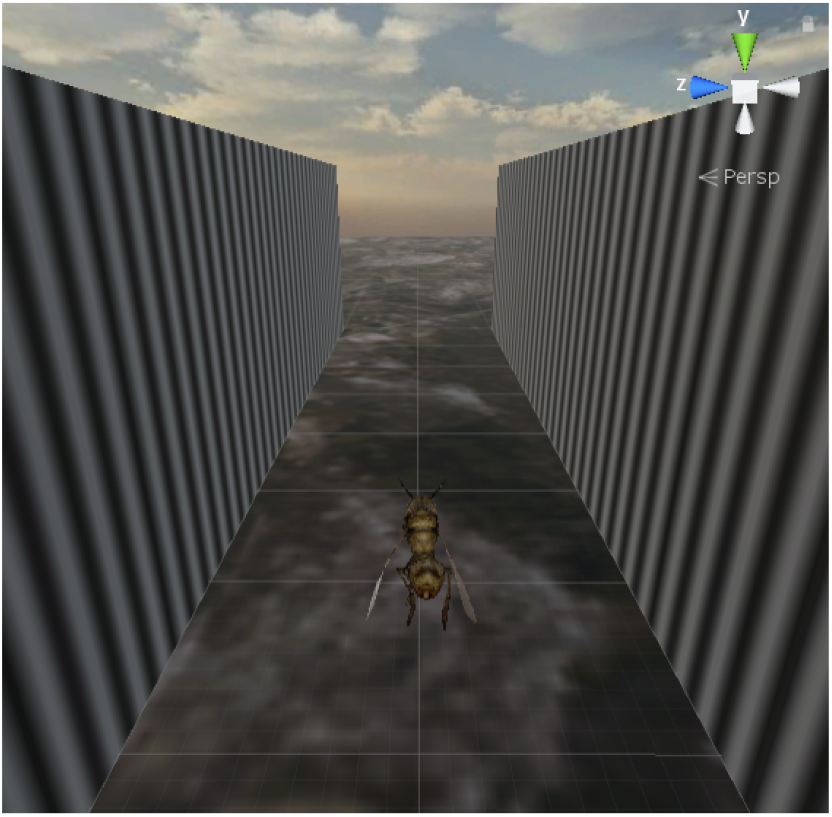
Unity simulation environment of the tunnel experiments. The virtual bee flies in a simulated tunnel with sinusoidal patterns on both walls. A demo video can be found at https://youtu.be/QXl95E71cTE.

In real tunnel behavioural experiments, honeybees can fly in the central of patterned tunnel even when the walls carry gratings of different spatial frequencies or contrasts [1]. Biologists suggest that honeybees can estimate the background speed independent of the spatial frequency and contrast, and adjust its position by balancing the angular velocity sensed by two compound eyes. The proposed AVDM possesses similar characteristics and demonstrates great potential in reproducing the honeybee’s tunnel centring responses. If the simulations work well, then it can verify the effectiveness and practicability of our proposed model.

In the first kind of simulations, the virtual bee starts at different positions in the patterned tunnel. We implement the AVDM into the two eyes of the virtual bee, and then test if the bee can perform a centring response as real bees. One of the simulation results with both walls carrying patterns of same spatial frequency (46 cycles m^−1^) is shown in Fig. 10. As can be seen, though the virtual honeybee is released at different start points, it can adjust its position using only visual information and finally fly along the central path of the tunnel. The result maintains the same if the spatial frequencies are changed (15, 20, 30, 40 cycles m^−1^) as long as the two walls carry the same pattern.

**Fig 10.**
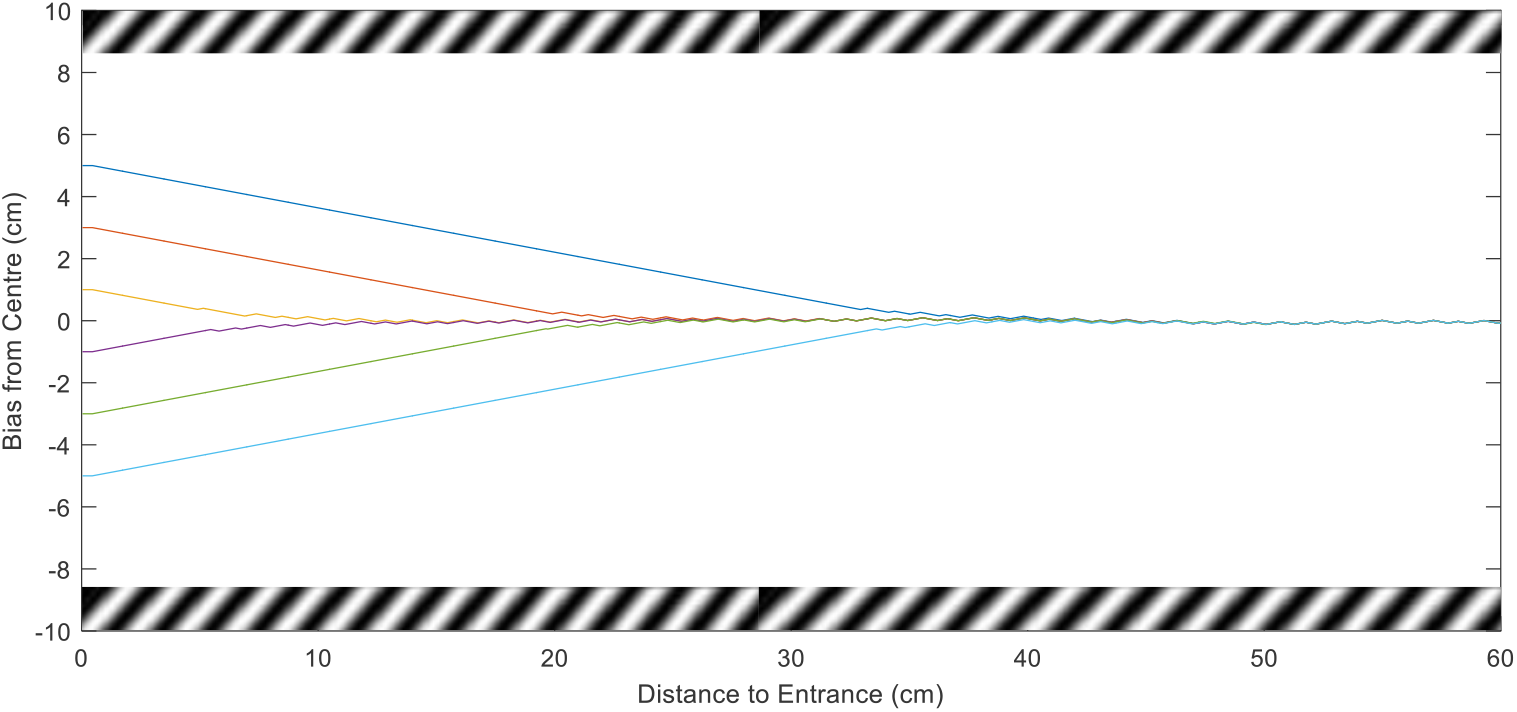
Tunnel centring from different start points. Routes of the virtual bee with AVDM implemented are recorded when they fly through the patterned tunnel from different start points. The flight paths are adjusted by the control scheme in Fig. 4.

Further, in the second kind of simulations, the spatial independence of the proposed model is investigated by changing the spatial frequency of one wall while keeping the other constant. The result is shown in Fig. 11. Though the spatial frequency of the right wall varies a lot, the virtual bee still flies through the tunnel with little biases from central path. This is in accordance with the biological experiment that the centring response is little affected by the spatial frequency of the pattern [34]. The biases may be caused by the difference of the estimated angular velocities when tested by different patterns (see Fig. 6). This means the model is not fully spatial independent. But real bumblebee behavioural experiments reveals that similar phenomenon can be observed in this situation [35], which indicates large rather than full spatial independence implemented in real bees’ neural system.

**Fig. 11.**
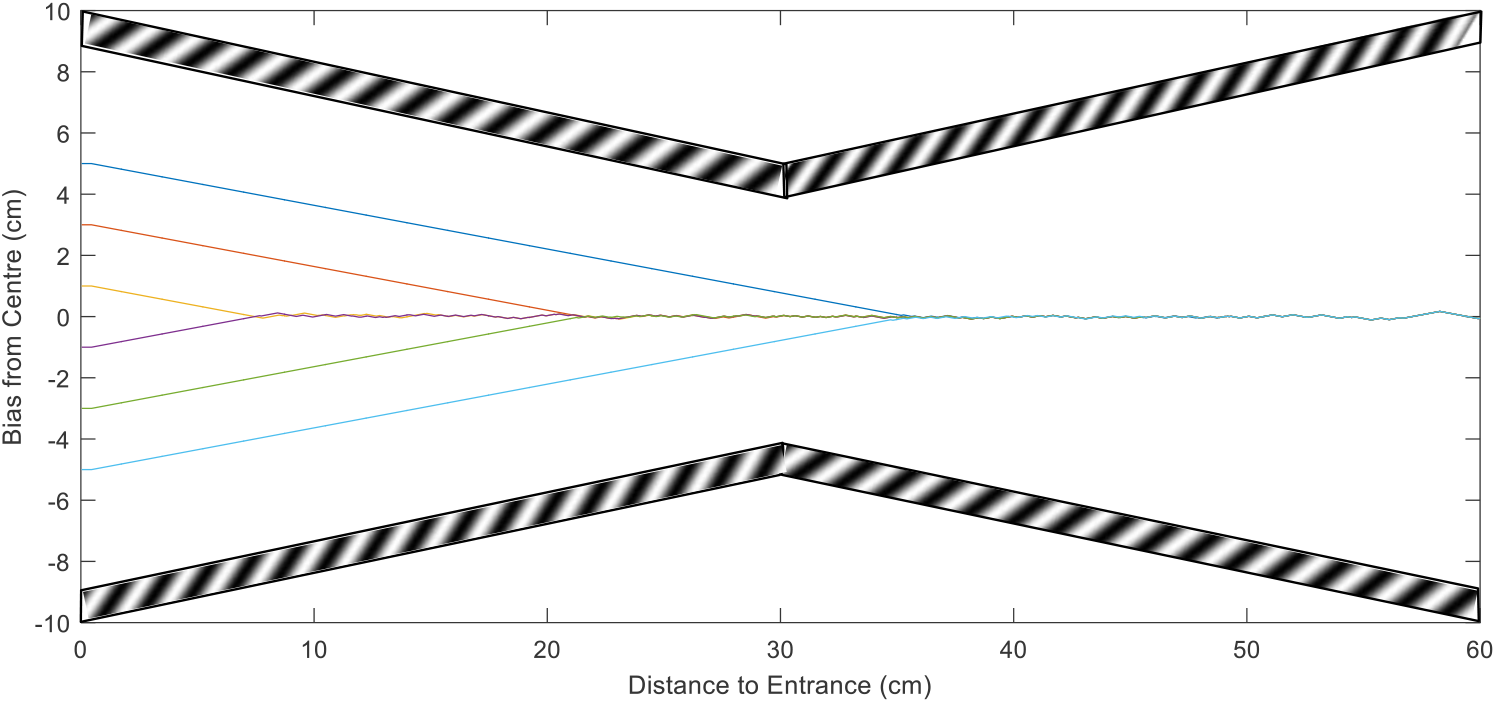
Tunnel centring with gratings of different spatial frequencies. The left wall is always carrying gratings of spatial frequency of 46 cycles m^−1^ while the the spatial frequency of right wall grating varies from 23, 35, 46, 69 cycles m^−1^. The trajectories of each simulation is recorded using different color lines.

In the third kind of simulations, the virtual bee is tested in a X-shape tunnel of which the width first decreases and then increases (see Fig. 12). Though released at different start points, the bee adjusts its lateral position well to fly towards the central path of the tunnel following the angular velocity balance strategy [36]. All these three kind of tunnel simulations reproduce similar behaviours of honeybees in patterned tunnel. However, this is not enough to show the proposed model estimate the angular velocity well. For example, a model capable of gauging distance towards wall using visual information can produce similar centring response too. A further simulation demonstrating that the AVDM do estimate the angular velocity rather than distance need to be designed.

**Fig. 12.**
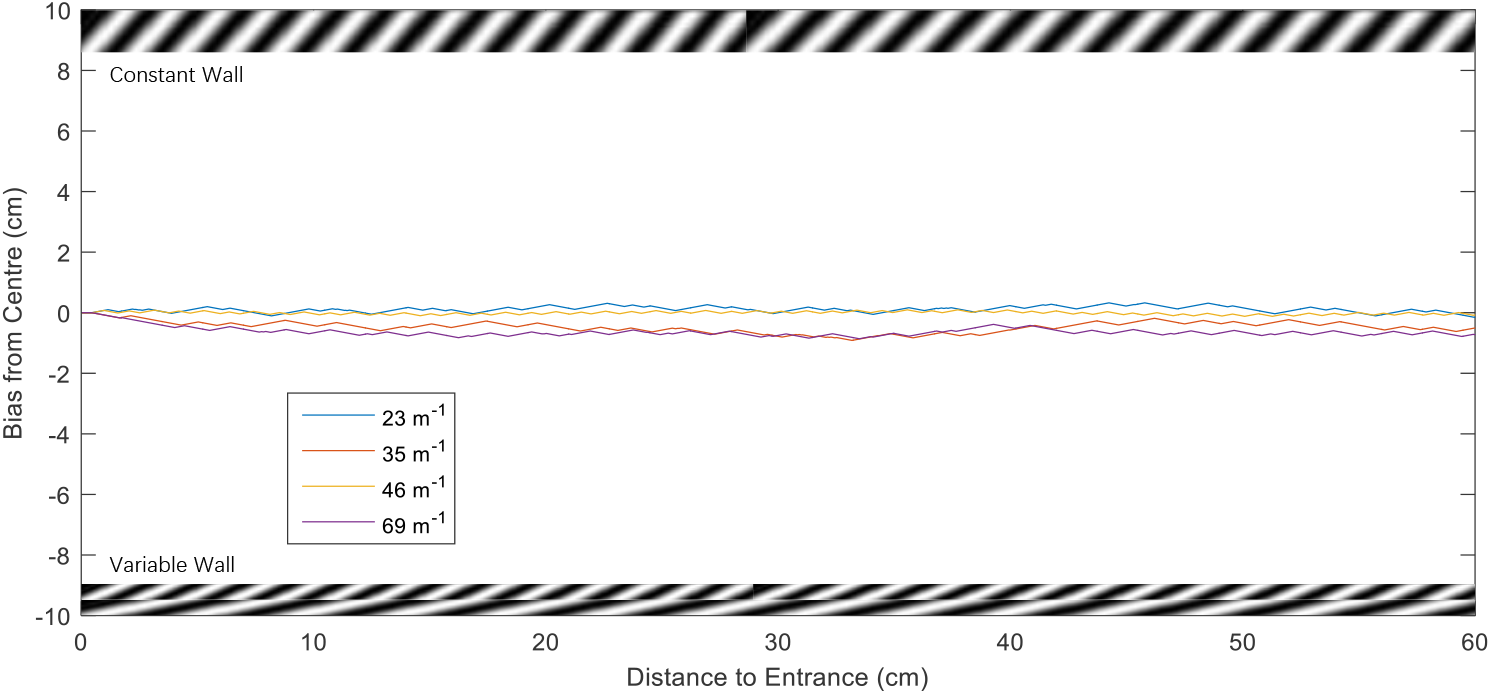
X-shape Tunnel centring simulations. The flight trajectories of the virtual bee starting from different points are shown in the same graph. The walls of the tunnel are covered with sinusoidal gratings of spatial frequency of 46 cycles m^−1^.

In tunnel experiments honeybees will move towards one wall if the wall moves along the same direction of their flights, and move towards the opposite side if the wall moves in the opposite direction [37]. The movement of the patterned wall affects the lateral position in tunnel. This indicates that honeybees do adjust positions by balancing the angular velocity on both eyes.

The proposed model is also tested in this scenario that one of the patterned wall moving along or against the flight direction. The simulation results are shown in Fig. 13. The virtual bee will move closer to left wall if the left wall is moving along the flight direction at a constant speed (much slower than the flight speed). This is because the angular velocity estimated on left eye is smaller than it was when the wall starts moving forward. Thereby the trajectory shifts towards to the moving wall to balance the angular velocities estimated on both eyes. On the contrary, the trajectory of the virtual bee shifts to right wall if the left wall is moving backward. Both coincide with the behavioural experiments of honeybees’ visual control [37], indicating that the proposed model can explain the tunnel centring behaviours of the honeybees very well.

**Fig. 13.**
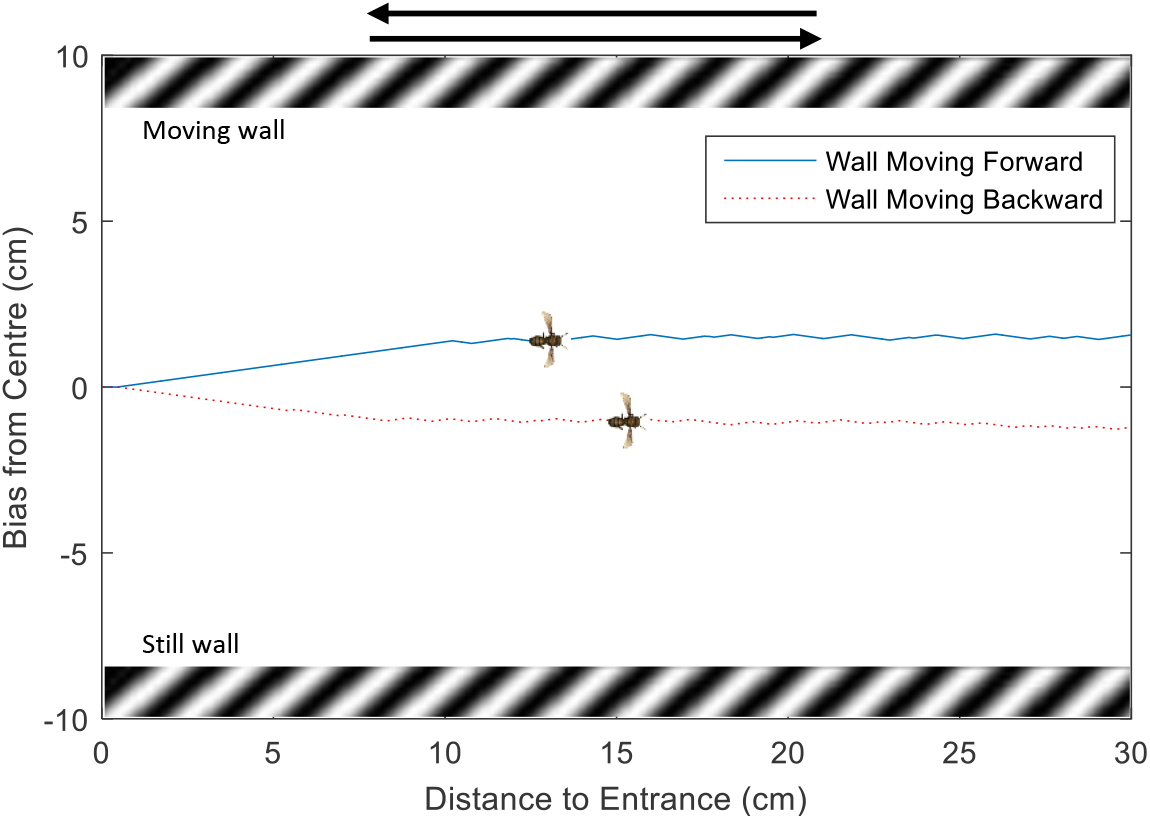
Tunnel centring with one wall moving forward and backward. The blue solid line indicates the flight trajectory when left wall is moving forward and the red dot line indicates the trajectory when left wall is moving backward.

### Terrain following simulation results

The accuracy of the angular velocity estimation has not been fully shown by the honeybees’ flight behaviours reproducing experiments in tunnel centring since the control scheme is triggered by the difference of the angular velocities estimated on two eyes. The error of the angular velocity estimation may be decreased by the substraction. To further verify the effectiveness of the proposed model, the AVDM is implemented in a virtual bee in terrain following simulations where the ground is covered with gratings (see Fig. 14). The performance of this visually guided flight task depends more on the angular velocity estimation accuracy, which providing an ideal chance to examine our proposed model. A series of terrain following simulations are designed. The flight trajectories and the ventral responses are recorded to see if the virtual bee can perform automatic terrain following using only visual information by estimating the angular velocity of image motion and regulating it to a constant value.

**Fig 14.**
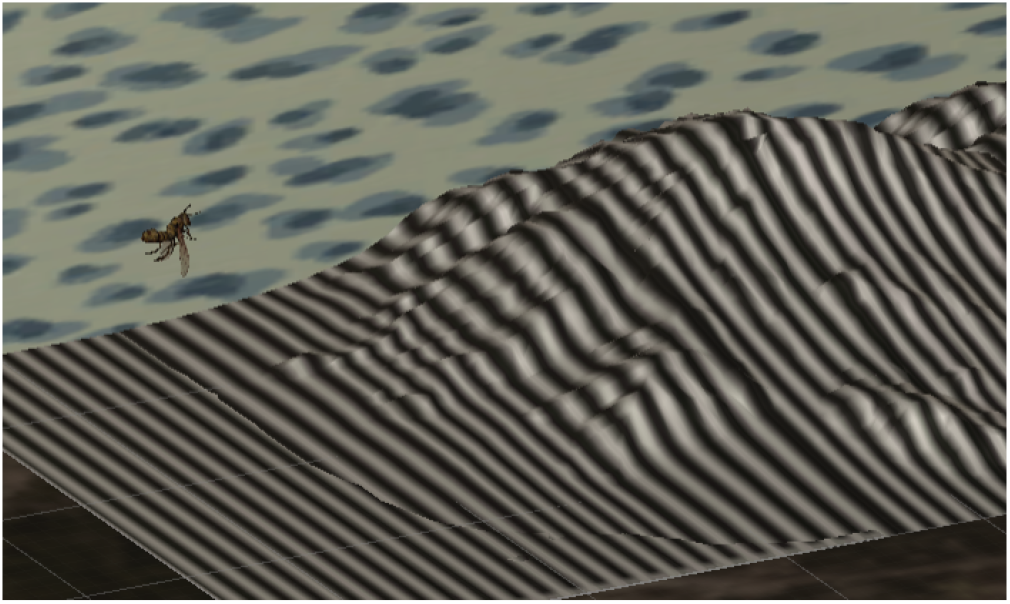
Unity simulation environment of the terrain following experiments. The bee flies over a textured terrain using only ventral visual information. A demo video can be found at https://youtu.be/jaYSuCJGAfc.

The virtual bee with AVDM implemented is first tested on a regular terrain covered with sinusoidal gratings. The virtual bee is released around a given height (25 cm) at a certain forward speed (50 cm/s). In the beginning phase, the bee is set to fly forward without changing its altitude (by setting the vertical lift equals its gravity). Then the preset angular velocity value is estimated using AVDM after the first few frames. By regulating the consecutive angular velocities to this preset value using the control scheme described in previous section (Fig. 5), we hope that the flight altitude will be adjusted automatically using only visual information. The result is shown in Fig. 15.

**Fig 15.**
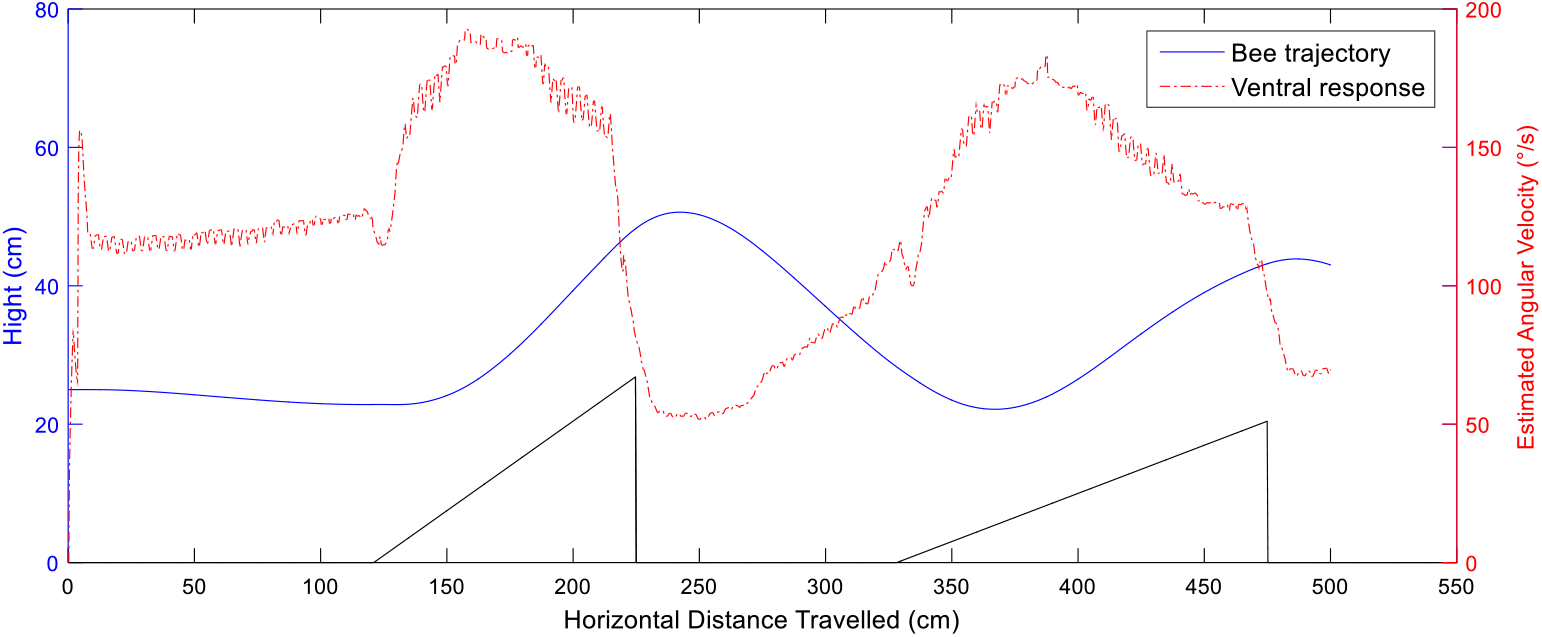
Terrain following flight trajectory and estimated angular velocities. The bee flight trajectory is recorded when flying over a terrain (black line) with sinusoidal gratings (30 cycles m^−1^), and the angular velocities estimated by the ventral eye are also shown indicating how the trajectory is affected by angular velocity regulating.

As can be seen, the angular velocity estimated by ventral camera is accurate and relatively keeps unchanged when the bee flies over flat terrain except first few frames. The ventral response increases when the bee flies closer to the undulating terrain, and vice versa. By changing the vertical lift according to the difference between estimated angular velocities and preset value, the virtual bee always keeps a distance from textured ground. Neither the flight speed nor the flight altitude is necessary to perform this visually guided task. In general, the proposed AVDM works well to navigate the bee flying over the patterned regular terrain. Similar terrain following experiments have been performed using aircrafts [14, 15]. The main difference is that they use the EMD sensor’s output rather than angular velocity to regulate the flight course. Our proposed model directly estimated the angular velocity to improve the accuracy of the regulation in terrain following.

To inspect the robustness of the terrain following under varies conditions, a series of controlled trials have been performed. First, the virtual bee is released at different initial heights, the terrain following trajectories are shown in Fig. 16(a). As can be seen, the virtual bee follows the terrain well maintaining certain distances according to the initial heights above the ground. The initial angular velocity presetting ensures the bee following the terrain well with dynamic initial heights. The lower the initial height is, the better the flight trajectory follows the terrain. When the initial height is 50 cm, the ventral angular velocity varies less since the undulation of terrain is relatively smaller at this height than other lower heights.

**Fig 16.**
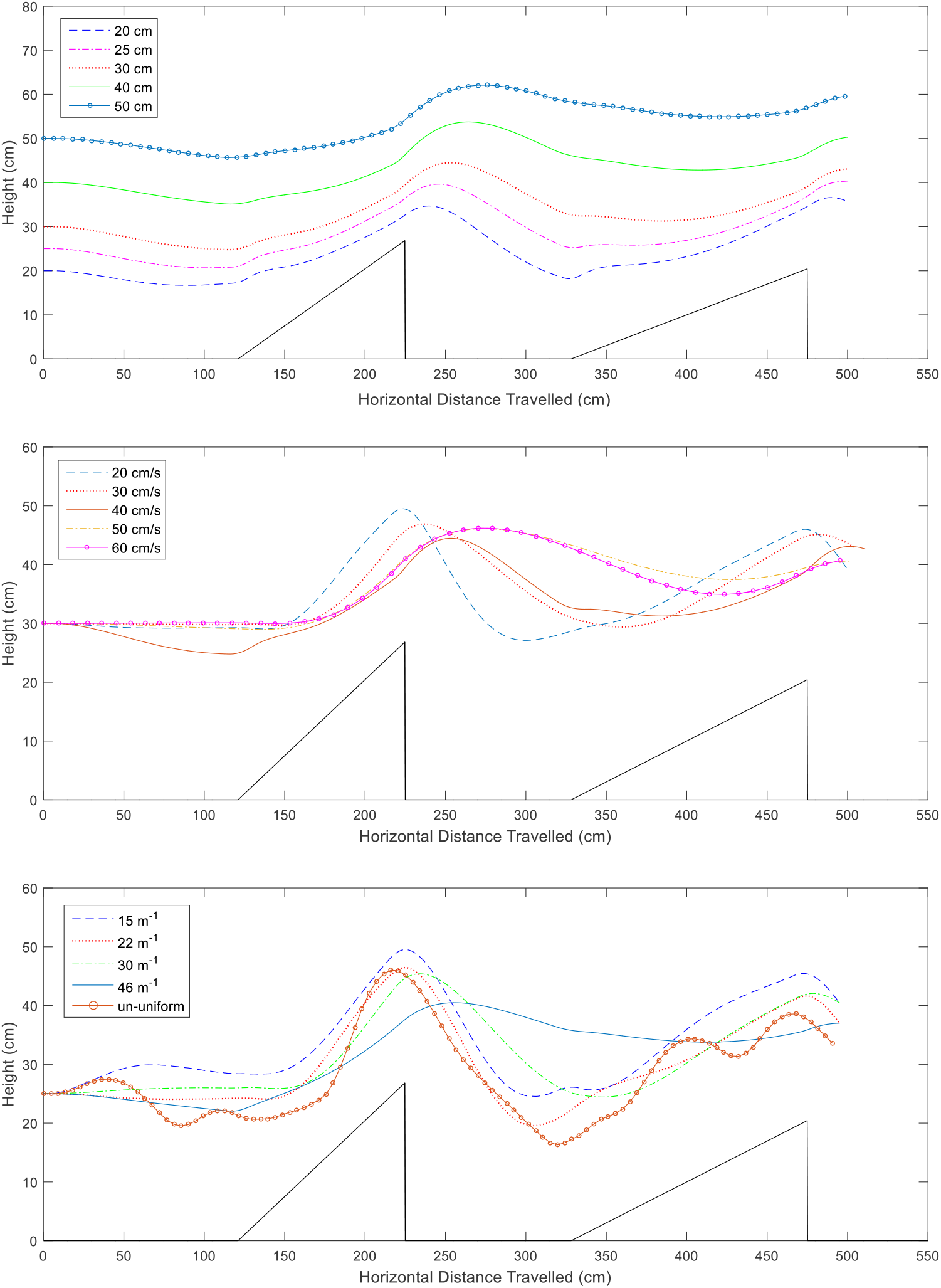
Controlled trials showing the robustness under different flight heights, speed and terrain gratings. (a) The virtual bee is released at different initial heights with a speed of 50 cm/s to fly over the terrain with sinusoidal grating (30 cycles m^−1^). (b) The virtual bee is released with different flight speeds at the height of 30 cm to fly over the terrain with sinusoidal grating (30 cycles m^−1^). (c) The virtual bee is released with a flight speed of 40 cm/s at the height of 25 cm to fly over the terrain with different gratings.

In second kind of controlled trials, the influence of the flight speed towards the terrain following is investigated. The flight speed is chosen from 20 cm/s to 60 cm/s, 39 and the trajectories are recorded in Fig. 16(b). The trajectories of all tested speeds follow the terrain well without any danger of crashing to the terrain using only visual information. The differences among trajectories are mainly caused by the control scheme. 39 The trajectory follows the terrain very well if the speed is 20 cm/s. It is harder to adjust the flight height smoothly if the virtual bee flies faster. A better control scheme may improve the performance of the following at varies speeds.

At last, the spatial independence of the proposed model in terrain following is inspected by covering the terrain with different gratings (see Fig. 16(c)). A wide range of spatial periods of gratings are chosen. All flight trajectories follow the terrain well given the same initial flight height and flight speed. Especially from the trajectory of the bee flying over gratings of non-uniform spatial frequency, we can tell spatial frequency do affect the terrain following a little. Again it fulfills our expectation of large rather than full spatial independence of the proposed model.

In order to see whether the model is stable under more complex scenarios, we also tested the virtual bee above a mountain shape terrain with irregular undulation covered with sinusoidal gratings. The result is shown in Fig. 17. The flight trajectories of the bee undulate above the mountain terrain and the flight altitude changes automatically according to the distance to ground by regulating the angular velocity. And whenever the bee gets closer to the ground, the increasing angular velocity estimated by the AVDM will trigger the controller to provide a higher vertical lift to help the bee get away from ground. This further verifies the effectiveness of the proposed AVDM in the terrain following simulations, showing the potential of applying in UAV’s (unmanned aerial vehicle) flight control.

**Fig 17.**
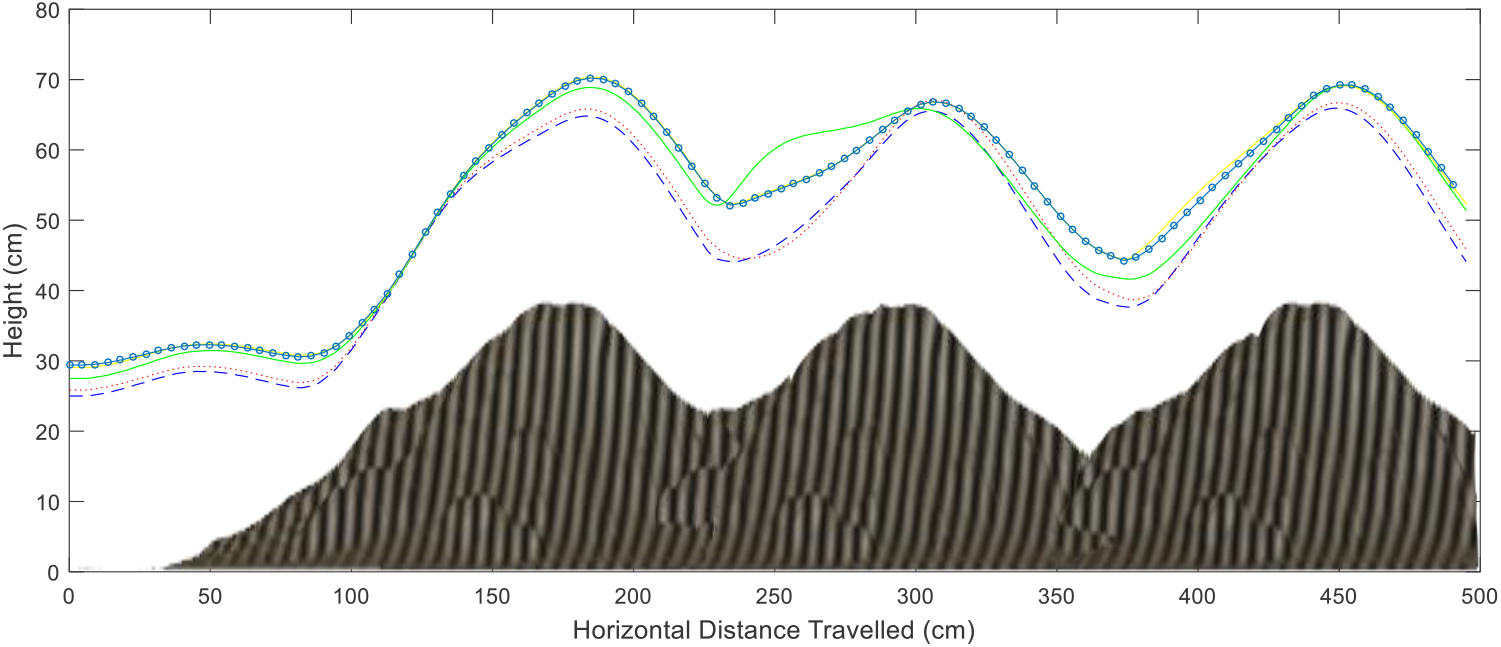
Multiple bee flight trajectories of terrain following over the mountain shape terrain. The successful terrain following flights over pattered mountain using only visual information show the robustness of the model and control scheme.

## Conclusion and Discussion

We have presented an angular velocity decoding model (AVDM), which estimates the visual motion speed combining both texture and temporal information from input signals. Before describing the model structure in detail, we have investigated the spatial and temporal resolutions of honeybees. This helps us to get a bio-plausible parameter setting which is very important for explaining honeybees’ flight behaviours. Then the model with three parts: elementary motion detection circuits, wide-field texture estimation pathway and angular velocity decoding layer is given. The model estimates the angular velocity very well with improved spatial frequency independence compared to the state-of-the-art angular velocity detecting models, when firstly tested by moving sinusoidal gratings. This spatial independence is vital to account for the honeybee’s tunnel centring response.

In order to investigate whether the model can account for observations of tunnel centring behaviours of honeybees, the model has been implemented in a virtual bee simulated by the game engine Unity, and a series of experiments have been designed. The simulation results show that the virtual bee can adjust its position to fly through the patterned tunnel by balancing the angular velocities estimated on both eyes under several circumstances. All tunnel stimulations reproduce similar behaviours of real bees. These are: flying towards central path in parallel tunnel and X-shape tunnel, little biases from central path when walls carrying gratings of different spatial frequencies and adjusting its position when one of the wall is moving forward or backward. This indicates that our model does provide a possible explanation of how honeybees estimate the image motion velocity and regulate the flight course in tunnel.

To further verify if the model can estimate the angular velocity of image motion accurately, the visually guided terrain following simulations have been carried out with a closed-loop control scheme to restore a preset angular velocity during the flight. The simulation results show that the virtual bee successfully flies over the undulating terrain under several settings using only visual information. This verifies the feasibility and robustness of the AVDM performing in various application scenarios, which gives further support for explaining honeybees’ visual motion detection and shows its potential in applications of unmanned aerial vehicles.

What’s more, the proposed model has potentials to simulate more behaviours of honeybees. For example, keeping constant angular velocity is also used in honeybees’ wall following behaviour [3, 38]. In addition, integrating the angular velocity decoded can provide the information of odometry. Since the proposed model estimates the angular velocity accurately, it helps to explain how honeybees gauge the flight distance [6].

Though the proposed model is designed primarily for estimating angular velocity under sinusoidal gratings, it can be easily generalized to deal with more patterns. In fact, even without any modification, the AVDM works well to decode angular velocity under patterns with clear edges such as square-wave grating and checkerboard pattern (data not shown). The model is further tested by virtual bee to follow an irregular snow mountain terrain [24]. The flight trajectory does not follow the terrain well as it does when tested by mountain covered with sinusoidal gratings. However, trajectories of successfully flying over the snow mountain without crushing indicate the potential of the model to navigate well in cluttered environment, see also **S2 Video**.

As for the model itself, we only provide motion detectors for the progressive and regressive directions. Motion detectors for upward and downward can be constructed similarly to form a more complete visual detection system for dealing with more complex and dynamic visual scenes. And the proposed model will be implemented into UAVs and to be tested in a real environment to verify its effectiveness in the near future.

## Supporting information

**S1 Video. A demo video for unity simulation of tunnel centring experiments.** Using Unity engine, the images received by two eyes can be processed separately in real time to estimate the image motion angular velocities on both sides. By balancing the angular velocities, the virtual bee shows similar behaviours like real honeybees. Simulation 1 shows the centering behaviour at different start points. And the simulation 2 shows the trajectory shifts when the wall of the tunnel moves forward and backward. The demo video can be found at https://youtu.be/QXl95E71cTE.

**S2 Video. A demo video for unity simulation of terrain following experiments.** Using Unity engine, the images received by ventral camera can be processed in real time to give an estimation of the image motion angular velocity. By regulating the angular velocity to a constant value, the virtual bee successfully accomplished three terrain following simulations. Simulation 1, 2, 3 shows the terrain following of the regular patterned ground, patterned mountain and irregular snow mountain. The demo video can be found at https://youtu.be/jaYSuCJGAfc.

## Author Contributions

Conceived and designed the experiments: HTW SY. Performed the experiments: HTW QF. Analyzed the data: HTW JP. Wrote and revised the paper: HTW QF HXW PB JP SY.

## Acknowledgments

This research is funded by the EU HORIZON 2020 project, STEP2DYNA (grant agreement No. 691154) and ULTRACEPT (grant agreement No. 778062); the National Natural Science Foundation of China (grant agreement No. 11771347)

